# Tillering structures the genetic variability of wheat vegetative growth and its plasticity under water deficit

**DOI:** 10.1101/2023.07.26.550706

**Authors:** Stéphane Leveau, Boris Parent, Francesco Giunta, Nathalie Luchaire, Llorenç Cabrera-Bosquet, Katia Beauchêne, Stéphane Jezequel, Rosella Motzo, Pierre Martre

## Abstract

Leaf expansion under drought drives the trade-off between water saving for later grain production and canopy photosynthesis. Fine-tuning leaf expansion could therefore become a target of genetic progress for drought-prone environments. However, its components (branching, leaf production and elongation) may have their own genetic variability and plasticity under drought, making hard to calibrate crop simulation models and specify breeding targets. In this study, we focused on the genetic diversity of bread wheat and durum wheat to determine the links and trade-offs between the underlying processes of leaf growth under drought and how it translates to leaf expansion at the whole plant and canopy level. For that, we used non-destructive imaging both in the field and controlled condition platforms to determine the dynamics of the components of shoot expansion and analyze their relative contribution to the genetic variability of whole-plant shoot expansion under drought. Results show that leaf expansion measured at plant level in controlled environment was associated with that measured at canopy level in the field, indicating that controlled phenotyping platforms can capture the genetic variability of growth in the field. Both whole-plant and canopy expansion were associated with tillering rate. In addition, the sensitivity of shoot growth and tillering to soil water deficit were correlated, indicating that both tillering ability and sensitivity to water deficit drive the genetic variability of shoot expansion. Overall, dissecting leaf expansion dynamics allowed determining the links between shoot expansion traits under drought, and provides key targets in phenotyping, modelling and breeding for drought environments.

## Introduction

The control of leaf area under drought and its dual consequences on photosynthesis and water saving is a major player in Genotype x Environment (G x E) interactions (Correia et al., 2022; Dhakar et al., 2023). Fine-tuning leaf expansion for drought envirotypes could therefore become a major target for genetic improvement in drought-prone environments (Tardieu et al., 2018). But the use of traits associated with leaf expansion as genetic levers in plant breeding may not be straightforward due to the trade-off between water saving traits (Faralli et al., 2019; Hatfield and Dold, 2019), positives for later stages of flowering and grain filling, but resulting in photosynthesis reduction. Thus, the best variety observed in one environment may be a very poor choice in a different drought scenario (Tardieu et al., 2018).

“Tuning growth to the environmental demand” (Rymen and Sugimoto, 2012) requires focusing not only on the plants constitutive growth potential but also on its ability to modulate its growth under drought (plasticity). Plasticity of plant architectural traits allows plants to modify their architecture when environment changes (Pierik et al., 2021). In the case of expansive growth, such phenotypic plasticity occurs at different levels of organization (Belaygue et al., 1996). Indeed, the whole-plant leaf area is the consequence of three underlying processes (branching or tillering, individual-leaf expansion rate, and leaf appearance rate). Such stress-adaptive plasticity of morpho-physiological processes has shown large genetic variability within wheat species and in wheat relatives (Suneja et al., 2019; Leveau et al., 2021). The fact that these processes have their own timing within the crop cycle and potentially their own genetic variability and response to water deficit makes it complex to analyze the genetic variability of whole plant growth under fluctuating environments. Indeed, the consequence is that there is not a unique response to water deficit, but potentially as many as there are water deficit scenarios (duration, intensity, timing; Koch et al., 2019) combined with genetic variability for the three processes underlying whole plant leaf expansion:

-The rate of leaf appearance is often considered to be stable and independent of the environment (except for photoperiod and temperature). In the only paper to our knowledge that considered the relative sensitivities of the underlying components of shoot expansion (in white clover; (Belaygue et al., 1996) found that leaf appearance rate was the least sensitive process. However, in wheat, Baumont et al. (2019) showed that the rate of leaf appearance is sensitive to the carbon status of the plant, and thus to the environmental conditions driving this status (intercepted radiation, photoperiod, atmospheric CO_2_ concentration, …), and that genetic variability exists for this trait. Overall, the relative contribution of leaf appearance rate to the genetic variability of whole-plant expansive growth under drought remains largely unknown.

-The response of individual-leaf expansion to soil water potential displays a large genetic variability in most studied species (e.g. in rice, Parent et al., 2010; and maize, Welcker et al., 2011), as its response to other environmental conditions in wheat (e.g. VPD; Leveau et al., 2021). In non-tillering maize hybrids, the response of individual leaves to soil water potential (SWP) correlated to that of whole plant (Lacube et al., 2020). However, its relative influence in tillering species such as wheat species has been far less studied.

-Tillering traits under drought have long been a pre-breeding target, especially for drought-prone environment (Richards, 1988; Wang et al., 2016). However, most studies have focused on the intrinsic value of tillering potential, with the aim of reducing tillering. Indeed, the advantage of reduced-tillering traits is close to a “drought avoidance strategy”, reducing leaf area in order to shift soil water availability from pre- to post-anthesis when periods of drought stress are frequent (Berry et al., 2003). A tiller inhibition gene (Atsmon and Jacobs, 1977) increases grain yield in dryland environments but not under wetter conditions (Hendriks et al., 2016). While it appears that tillering is a key process for phenotypic plasticity (Sanad et al., 2016), we could find only few results on the plasticity of free-tillering genotypes and the associated genetic diversity, scatters into more general studies.

Field networks cannot explore the impact of each combination of plasticity-related traits across the full range of drought scenarios. Model-based ideotyping (Senapati et al., 2022) uses a crop model with genetic parameters, capable of simulating the Genotype x Environment x Management interactions under a wide variety of present and future local scenarios (Parent et al., 2018). In the case of whole-plant shoot expansion, if its underlying components and their sensitivities to environmental constraints were fully independent, with different genetic controls, this would imply a large number of genetic parameters impossible to measure in hundreds of genotypes. However, the independency of these traits can be questioned because some common underlying processes can drive several components, such as tissue water potential and turgor, and the potential rate of one process (e.g., individual leaf expansion rate) is often linked to its sensitivity to water deficit, the largest plants being the most sensitive to water stress (as in maize; Welcker et al., 2011).

Determining such parameters for hundreds of genotypes requires high-throughput phenotyping (HTP) techniques. Such HTP platforms (Tardieu et al., 2017; Janni and Pieruschka, 2022) have been developed, in both controlled environment (Cabrera-Bosquet et al., 2016) or in the semi-controlled field (Beauchêne et al., 2019). In this landscape, HTP platforms under controlled conditions allow a fine characterization of a number of traits in thousands of individuals together with a fine control of environmental conditions, using consistent and reproducible methods between experiments. In parallel, field platforms provide crop growth conditions that are closer to agronomic conditions, but they rarely allow a fine control of environmental conditions such as well-defined water deficit scenarios. We should therefore take advantage of their complementarity to better understand and characterize the diversity of physiological traits associated with growth plasticity.

In this study, we aimed at determining the links between the components of shoot expansion under water deficit, and their relative contribution to the genetic variability of whole plant and canopy expansion in bread wheat and durum wheat. For that, we used a non-destructive imaging platform, (Cabrera-Bosquet et al., 2016), combining automatic measurements of shoot area over time (> 25,000) with manual measurements of individual leaf length and width, tiller number, and leaf number over time (> 10,000 measurements for each trait), to analyze the dynamics of whole-plant shoot growth and those of three underlying processes (tillering, leaf production, and leaf elongation). Results show that the genetic variability of whole-plant shoot growth followed that of tillering. Because this result was obtained in a phenotyping platform which may have exacerbated the impact of tillering, we then questioned how such result could be translated into the field by analyzing the same genotypes in a field platform equipped with a self-guided imaging rover (Madec et al., 2017). Because growth rates measured in both the platform and in the field were highly correlated, we analyzed the sensitivities of growth components in the platform only, by using well-defined drought scenarios. Our results demonstrate that the observed diversity of plasticity for shoot area expansion is mostly driven by that of tillering.

Overall, this work can have large consequences in (i) phenotyping, by demonstrating that the use of a phenotyping platform under controlled conditions can capture the genetic variability of growth rates measured in the field, (i) in modelling, by describing the phenotypic space of shoot expansion traits, therefore, simplifying the parameterization of the plasticity of leaf area under drought in crop growth models, and (iii) in breeding, by determining the genetic variability for target traits under drought in two species.

## Results

### Dynamic responses of leaf area expansion and underlying processes to water deficit

The dynamics of whole-plant leaf area (shoot area, SA) expansion was analyzed for 44 genotypes of bread wheat and 33 genotypes of durum wheat under well-watered scenario (S1) and three water-deficit scenarios contrasting in their timing and duration (S2 to S4, see Material and Methods for details) in the *PhenoArch* phenotyping platform (Cabrera-Bosquet et al., 2016). These genotypes were selected for presenting contrasted sensitivities under non-optimal conditions in the field (Material and Methods and Supplemental Fig. S1). Time courses of SA over thermal time after leaf-3 emergence were fitted with a cubic smoothing spline function (Fig. 1A). For plants under well-watered conditions (scenario S1), time courses of SA showed a typical sigmoid curve with a period of exponential increase, followed by an almost-linear increase, a period of slowing-down, a maximum value, and a decrease due to leaf senescence after anthesis.

**Figure 1.**
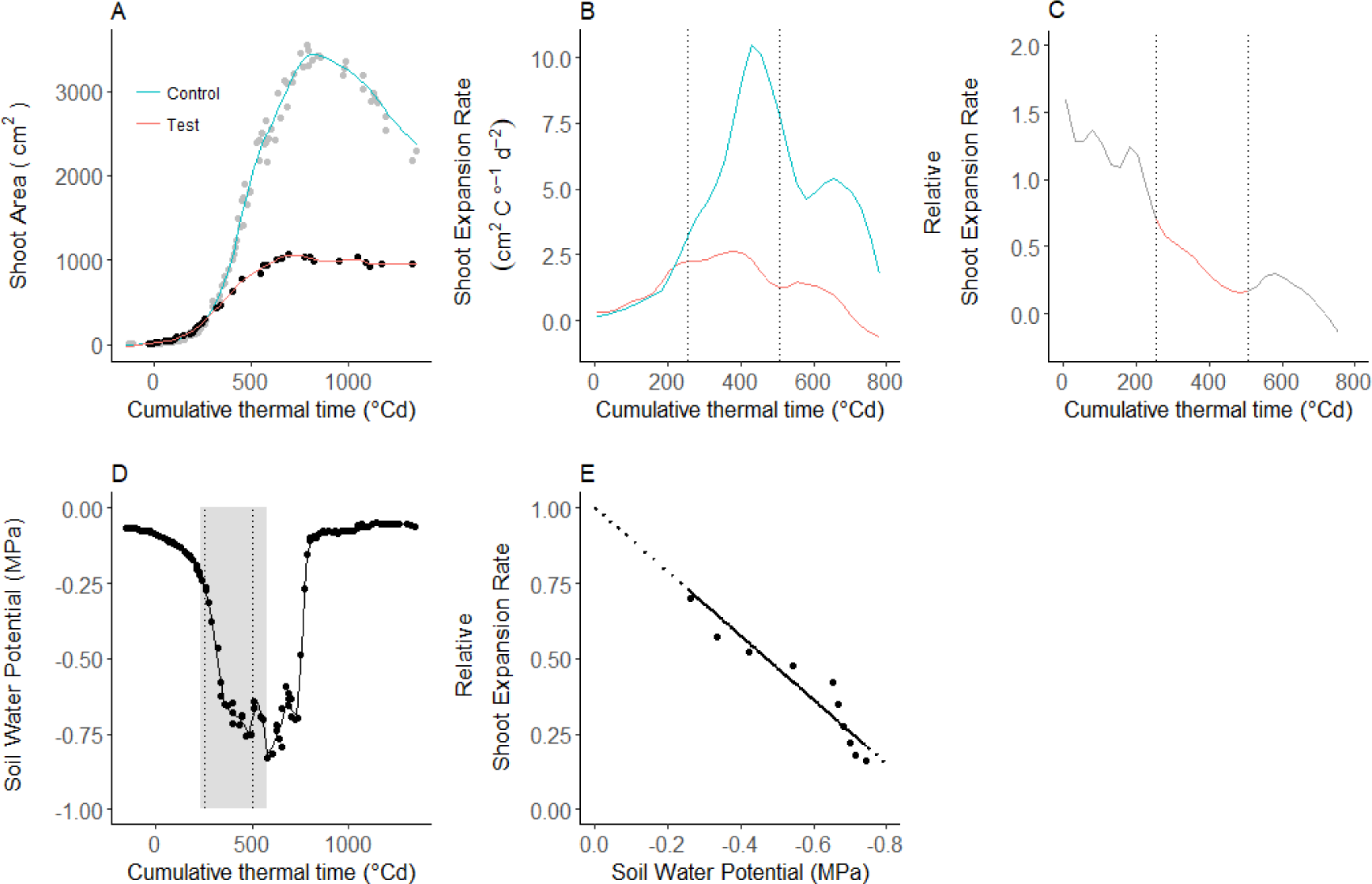
Example of calculation of dynamic variables and their sensitivity to soil water potential. Whole-plant leaf area (shoot area, SA) for the durum wheat cultivar Balsamo in the well-watered (control, scenario 1) and the water deficit scenario S4 (long water deficit). Thermal time was calculated from leaf 3 appearance. (A) Dynamics of SA. A cubic smoothing spline (lines) was fitted to experimental data (circles). In the control treatment, the spline function was fitted to the data of all plants (*n* = 3 to 6 independent replicates depending on the group), while for the water deficit scenario, data for each plant were fitted separately (one plant only is shown). (B) Dynamics of whole-plant leaf area expansion rate (shoot expansion rate, SER. SER was calculated as the first derivative of the spline function equation fitted to SA. The vertical dashed lines indicate the period beginning and ending when SER in the control scenario was 30% and 20% of its maximum value, respectively (also shown in C and D). (C) Dynamics of relative SER. Relative SER was calculated by dividing SER of each plant in the water-deficit scenario (red curve in B) by that of plants in the well-watered scenario (blue curve in B). (D) Dynamics of Soil Water Potential (SWP) for the plant under water deficit displayed in red in panels A,B,C. The grey band indicates the period beginning when SWP in water deficit scenario reached −0.2 MPa and ended on the day the plants were first rewatered. (E) relative SER versus SWP during the consensus period defined by the vertical dashed lines and the gray band. Solid line is linear regression forced to 1 when SWP is null. The slope of this regression corresponds to the sensitivity of SER to SWP for this plant.

Whole-plant leaf expansion rate (shoot expansion rate, SER), calculated as the first derivation of SA curves displayed a bell shape (Fig. 1B), with maximum values between 400 and 800°Cd after leaf-3 appearance depending on the genotype. The three water deficit scenarios affected the time course of SER. In scenario S4 (a 31-d period of water deficit starting at the beginning of tillering), SER increased similarly to that of well-watered plants during a first period (Fig. 1B), and then increased more slowly from around 250°Cd after leaf-3 appearance, when SWP decreased below −0.2 MPa (Fig. 1D). It resulted that the relative value of SER of plants under water deficit compared to that of well-watered plants was close to one until around 200°Cd after leaf-3 appearance (Fig. 1C), when the soil water potential (SWP) was about −0.2 MPa (Fig. 1D), and then decreased almost linearly as SWP continued to decrease. When plotted against SWP, during a period combining maximum values of SER in well-watered conditions (S1) and between the time at which SWP decreased below −0.2 MPa and the day of first rewatering, the relative SER showed a linear response to SWP (Fig. 1E). All genotypes showed similar SER time course and response to SWP, but with different absolute values (Fig. 2 for six genotypes).

**Figure 2.**
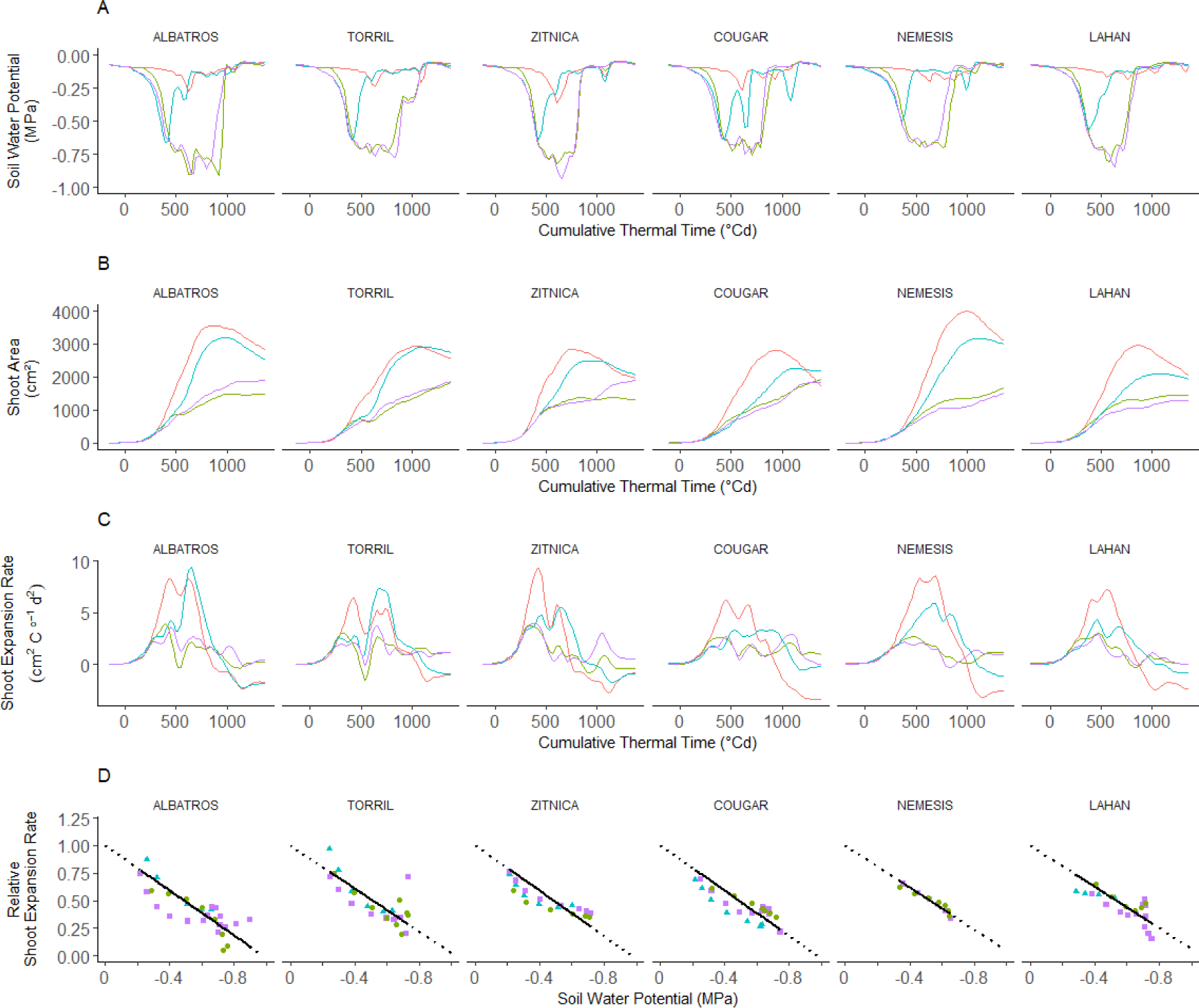
Time courses of whole-plant leaf area (shoot area) and its sensitivity to soil water potential for the six cultivars of group A (four bread-wheat cultivars: Albatros, Torril, Zitnica, and Cougar, and two durum wheat cultivars: Nemesis and Lahan) under the four watering scenarios. Thermal time was calculated from leaf 3 appearance. (A) to (C), dynamics of soil water potential, shoot area, and shoot expansion rate, respectively. (D) relative shoot expansion rate versus soil water potential. In (A) and (B) lines are cubic smoothing splines fitted to the experimental data. In (C) lines are the first derivatives of the fitted splines shown in (B). In (C) symbols are the calculated daily values during the period determined as illustrated in Figure 1, and lines are linear regressions forced to unit when soil water potential is null. Red lines, well-watered scenario (S1); blue triangles and lines, early drought scenario (S3); green circles and lines, late drought scenario (S2), and purple squares and lines, long-lasting drought scenario (S4).

Other drought scenarios affected SER in a similar way but with different timings (Fig. 2). The period defined above to calculate the sensitivity of SER to SWP resulted, for all genotypes, in a unique response in the three tested water-deficit scenarios (Fig. 2D).

Similar analyses of the time courses of absolute values, rates, relative rates and sensitivities to SWP were performed for the three underlying components of shoot area (i.e. number of appeared tillers, number of appeared leaves, and cumulative leaf length; Table 1). The tiller appearance rate (TiR) of well-watered plants (S1), calculated from the number of appeared tillers over time, had a time course very-similar to that of SER (Fig. 3, for one genotype and Supplemental Fig. S3, for six genotypes), but with maximum values occurring slightly sooner compared to SER. For plants under water deficit, TiR diverged from that of well-watered plants at around −0.2 MPa, similarly to SER. This resulted in a unique linear response of relative TiR to SWP in the three water-deficit scenarios for each genotype (Fig. 3 and Supplemental Fig. S3) but with higher sensitivity compared to that of SER.

**Table 1:** Processes and variables analyzed in the platform (A), and in the field (B).

| Process | Variables |  |  |  |  |
| --- | --- | --- | --- | --- | --- |
|  | Absolute values |  | Rates |  | Sensitivity to soil water potential |
|  | Dynamic variable (observed) | Metrics | Dynamic variable | Metrics |  |
| A. Phenotyping platform |  |  |  |  |  |
| Whole-plant shoot expansion | Shoot area | Maximum shoot area | Shoot expansion rate | Maximum shoot expansion rate | Sensitivity of shoot expansion rate |
|  | (SA, cm<sup>2</sup> plant<sup>-1</sup>) | (SA<sub>max</sub>, cm<sup>2</sup> plant<sup>-1</sup>) | (SER, cm<sup>2</sup> °Cd<sup>-1</sup>) | (SER<sub>max</sub>, cm<sup>2</sup> °Cd<sup>-1</sup>) | (S<sub>SER</sub>, MPa<sup>-1</sup>) |
| Leaf appearance | Leaf number on main stem | Final leaf number on main stem | Leaf appearance rate | Average leaf appearance rate | Sensitivity of leaf appearance rate |
|  | (LN, leaf main stem<sup>-1</sup>) | (FLN, leaf main stem<sup>-1</sup>) | (LAR, leaf °Cd<sup>-1</sup>) | (LAR<sub>mean</sub>, leaf °Cd<sup>-1</sup>) | (S<sub>LAR</sub>, MPa<sup>-1</sup>) |
| Tiller appearance | Tiller number | Maximum tiller number | Tiller appearance rate | Maximum tiller appearance rate | Sensitivity of tiller appearance rate |
|  | (TN, tiller plant<sup>-1</sup>) | (TN<sub>max</sub>, tiller plant<sup>-1</sup>) | (TiR, tiller °Cd<sup>-1</sup>) | (TiR<sub>max</sub>, tiller °Cd<sup>-1</sup>) | (S<sub>TiR</sub>, MPa<sup>-1</sup>) |
| Individual leaf elongation | Cumulative leaf length on main stem | Maximum cumulative leaf length | Leaf elongation rate | Maximum leaf elongation rate | Sensitivity of leaf elongation rate |
|  | (CLL<sub>max</sub>, cm plant<sup>-1</sup>) | (CLL<sub>max</sub>, cm plant<sup>-1</sup>) | (LER, cm °Cd<sup>-1</sup>) | (LER<sub>max</sub>, cm °Cd<sup>-1</sup>) | (S<sub>LER</sub>, MPa<sup>-1</sup>) |
| B. Field |  |  |  |  |  |
| Canopy expansion | Green Area Index | Maximum GAI | Rate of change of GAI | Maximal rate of change of GAI |  |
|  | (GAI, m<sup>2</sup> m<sup>-2</sup>) | (GAI<sub>max</sub>, m<sup>2</sup> m<sup>-2</sup>) | (GAIR, m<sup>2</sup> m<sup>-2</sup> °Cd<sup>-1</sup>) | (GAIR<sub>max</sub>, m<sup>2</sup> m<sup>-2</sup> °Cd<sup>-1</sup>) |  |

**Figure 3.**
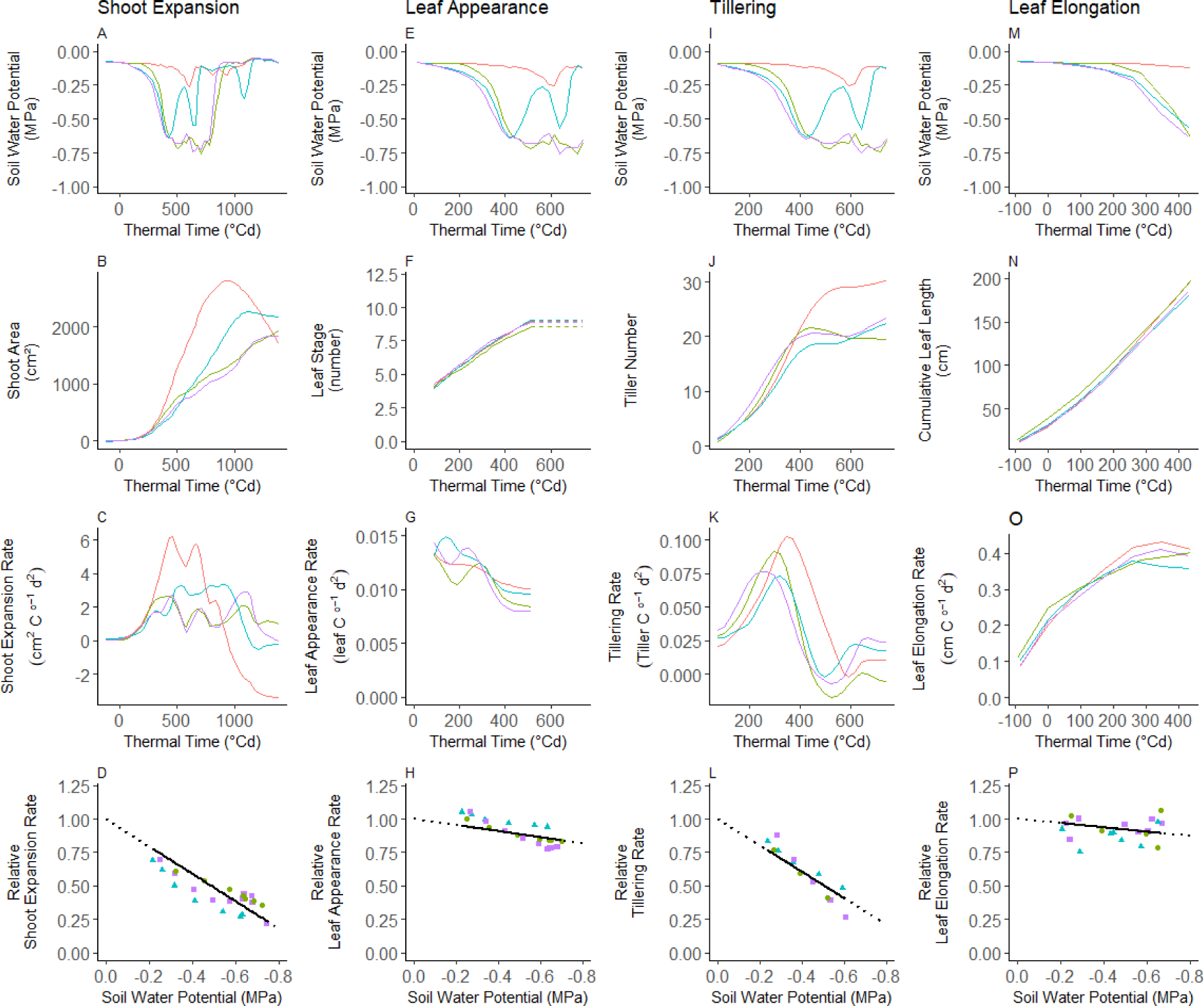
Time courses and sensitivities to soil water potential (SWP) of whole-plant leaf area (shoot area, **SA**) and its three underlying processes. Data are for the bread wheat cultivar Cougar grown under the four watering scenarios. Thermal time was calculated from leaf 3 appearance. (A) to (D), SA related variables calculated as in Figures 1 and 2. (E) to (H) Leaf appearance related variables. (I) to (L) Tillering related variables. (M) to (P) Leaf elongation related variables. The first row (A, E, I, and M) of panels shows SWP in the four treatments during the period of interest for these variables. The second row (B, F, J, and N) of panels shows cubic smoothing splines fitted to experimental data. The third row (C, G, K, and O) of panels shows rates as the first derivative of the fitted splines shown in the above row. The fourth row (D, H, L, and P) of panels shows the sensitivities of rates to SWP. Symbols are the calculated daily values during the period determined as illustrated in Figure 1, and lines are linear regressions forced to unit when SWP is null. Red lines, well-watered scenario (S1); blue triangles and lines, early drought scenario (S3); green circles and lines, late drought scenario (S2), and purple squares and lines, long-lasting drought scenario (S4). Figure 2 and Supplemental Figures 4, 5, and 6 show similar plots for shoot area, tiller appearance, main-stem leaf elongation and main-stem leaf appearance for the six genotypes of group A, respectively. Note that for each process the thermal time period was adjusted to show only the period of interest for this specific process.

Because water-deficit can affect the growth duration of individual organs (Verbraeken et al., 2021), we analyzed the rate of elongation of all growing leaves on the main stem together, to avoid a possible compensating effect by duration. Cumulated leaf length on the main stem over time for plants under water deficit was closer to that of well-watered plants than that was observed for SA and the number of appeared tillers (Fig. 3). This resulted in small differences in the elongation rate of individual main-stem leaves (LER) between the four scenarios (Fig. 3 and Supplemental Fig. S4), and a lower sensitivity of LER to SWP compared to that of TiR and SER.

In all scenarios and for all genotypes, leaf appearance rate (LAR) on the main stem calculated between leaf 3 and flag leaf appearance showed a slight decrease with time (Fig. 3 and Supplemental Fig. S5). As a result, the sensitivity of LAR to SWP was very low, even positive for some genotypes (Supplemental Tables S3 and S4).

### Under well-watered conditions, the genetic variability of tiller appearance rate is the main contributor to that of whole plant leaf expansion

To understand the relative contribution of the genetic variability of the three underlying processes to that of shoot expansion in well-watered conditions, we first compared the coefficients of variation of the maximum rates for shoot expansion (SER_max_), tiller appearance (TiR_max_), leaf appearance (LAR_mean_) and individual leaf elongation (LER_max_) in well-watered conditions (Fig. 4). For both species, the genetic variability of LAR_mean_ and LER_max_ was small compared to that of SER_max_, as indicated by their lower coefficients of variation. On the contrary, that of TiR_max_ was higher than that of SER_max_, especially for bread wheat.

**Figure 4.**
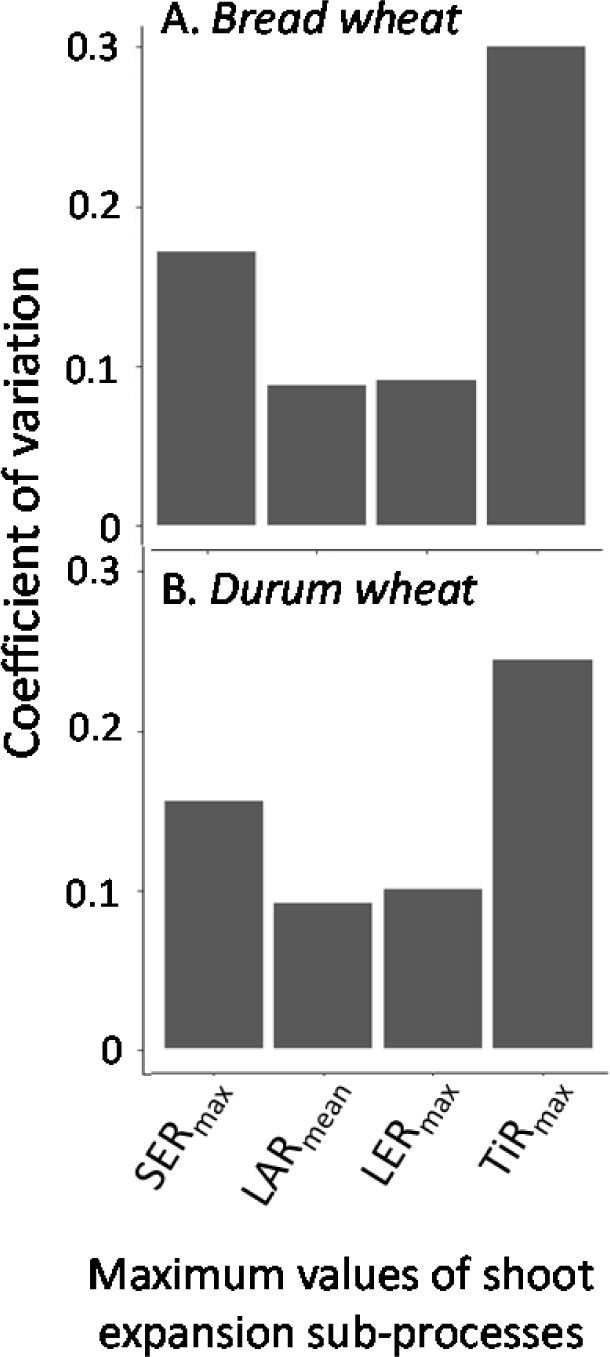
Coefficient of variation of maximum shoot expansion rate (SER_max_) and its three underlying processes, main-stem leaf appearance rate (LAR_mean_), maximum leaf elongation rate (LER_max_), and maximum tiller appearance rate (TiR_max_) for 44 bread wheat (A) and 38 durum wheat (B) cultivars grown under well-watered conditions (scenario 1) in the phenotyping platform.

An analysis of variance (Table 2A) showed that all three components explained a percentage of the variance of SER_max_, but with a higher total variation attributable to TiR_max_. This was confirmed by looking to the bivariate relationships between these variables (Fig. 5), which showed significant relationships between SER_max_ and TiR_max_, either in each species or considering the two species together. This was not the case for LAR_max_ and LER_max_ for which the correlation with SER_max_ was not significant. While less significant than for LAR_max_ and LER_max_, the final leaf number on the main stem (FLN) was associated with SER_max_ in the ANOVA. Indeed, the bivariate relationships showed that all components of growth were associated to some extent with FLN (Supplemental Fig. S6). This was especially the case for TiR_max_, which was positively associated with FLN (*r* =0.56, *P* < 1 × 10^-9^).

**Table 2.** Analysis of variance (ANOVA) of whole-plant shoot expansion processes (A: rates; B: sensitivities to soil water potential) as a function of their three underlying components, leaf appearance, tillering and leaf elongation. (A) ANOVA for the maximum value of shoot expansion rate (SER_max_) as a function of mean leaf appearance rate (LAR**_mean_**), maximum main-stem leaf elongation rate (LER**_max_**), and maximum tiller appearance rate (TIR**_max_**) and the potential covariable main-stem final leaf number (FLN). (B) ANOVA for the sensitivity of shoot expansion rate as a function of that of ssleaf appearance rate (*S*_LAR_), that of tiller appearance rate (*S*_TIR_), and that of leaf elongation rate (*S*_LER_).

|  | Degree of freedom | Sum of squares | Mean square | F-value | P-value |
| --- | --- | --- | --- | --- | --- |
| A. Predictor |  |  |  |  |  |
| $LAR_{mean}$ | 1 | 12.30 | 12.30 | 10.77 | < 0.01 |
| $TIR_{max}$ | 1 | 22.40 | 22.38 | 19.59 | < 0.001 |
| $LER_{max}$ | 1 | 17.88 | 17.88 | 15.68 | < 0.001 |
| FLN | 1 | 6.01 | 6.10 | 5.34 | < 0.05 |
| Residuals | 72 | 82.27 | 1.14 | NA | NA |
|  | Degree of freedom | Sum of squares | Mean square | F-value | P-value |
| B. Predictor |  |  |  |  |  |
| $S_{LAR}$ | 1 | 0.001 | 0.001 | 7.641 | <0.01 |
| $S_{TIR}$ | 1 | 0.004 | 0.004 | 32.476 | <0.001 |
| $S_{LER}$ | 1 | 0.001 | 0.001 | 9.052 | <0.01 |
| Residuals | 72 | 0.009 | 0 | NA | NA |

**Figure 5.**
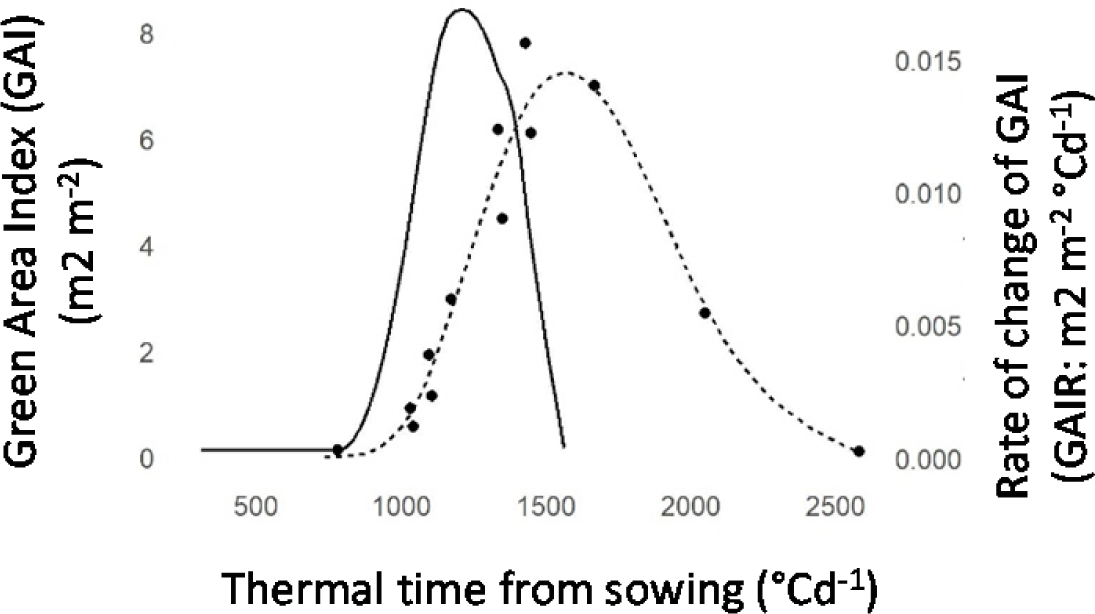
Time courses of absolute values and rate of change of green area index for the bread wheat genotype “2109.36” grown in the field at Gréoux-les-bains, France during the 2017-2018 growing season. Data are mean for *n* = 2 independent replicated. Dashed lines are cubic smoothing spline fitted to the data. Solid lines are the first derivate (rate) of the fitted splines.

Overall, tillering was the underlying process with the highest relative variability, and the most associated with shoot expansion, indicating a higher relative contribution to the genetic variability of shoot expansion, compared to leaf appearance and individual-leaf elongation.

### Tillering rate measured in the platform correlated with canopy expansion in the field

Because the above results were obtained on individual plants in the platform, we questioned if such results could be extrapolated at the canopy level in the field. For that, we analyzed the time courses of canopy expansion for the 44 bread wheat and 33 durum wheat cultivars, grown respectively during the 2017-2018 and 2018-2019 seasons. For both species, green area index (GAI) was determined with a self-guided rover (Madec et al., 2017) imaging plots every about 500°Cd during the whole crop growth season, and with a higher frequency (every 100°Cd) from 1000°Cd to 1500°C after sowing (Fig. 5).

GAI showed similar dynamics as those observed for SA in the platform, with a period with an almost-linear increase, followed by a period of decrease due to leaf senescence. Fitting spline curves to observed GAI data allowed us to calculate the rate of change (GAIR) as a function of thermal time (Fig. 5). The maximum values of GAIR (GAIR_max_) correlated with SER_max_ (Fig. 6) for bread wheat (*r* = 0.5; *P* < 0.01), and to a lower extend for durum wheat (*r* = 0.41, *P* = 0.09), indicating that growth rates measured on individual plants in the platform can be extrapolated to the field. Surprisingly, correlations found between TiR_max_ estimated in the platform and GAIR_max_ in the field were significant (Fig. 6) for both species (*P* < 0.05), indicating that the genetic variability of tillering potential measured in the platform reflected, to some extent, that of canopy expansion in the field.

**Figure 6.**
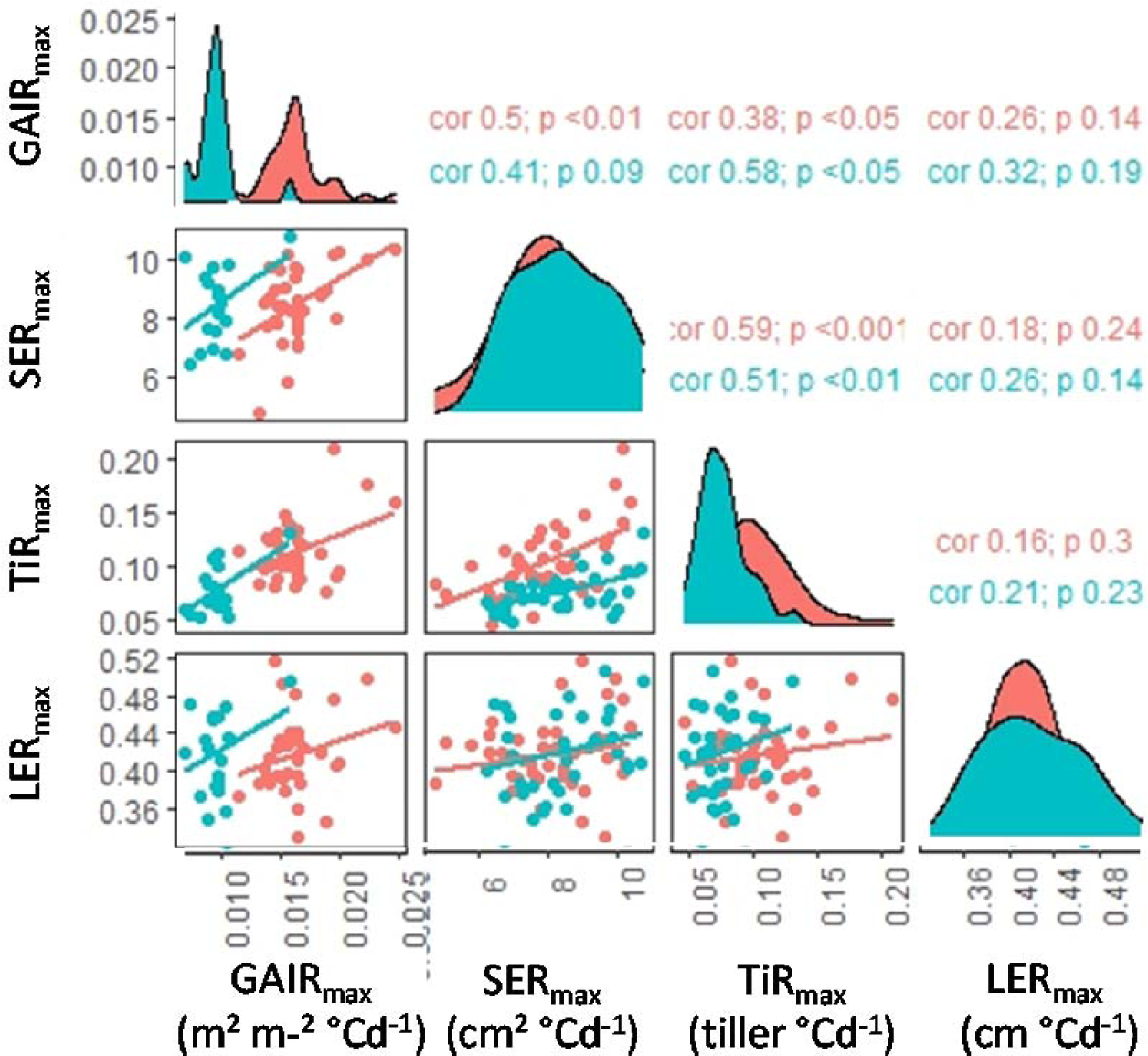
Scatter plot matrix of bivariate relationships between variables linked to shoot growth measured in the field or in the platform for all genotypes of bread wheat (red) or durum wheat (green). GAIR_max_, maximum rate of change of green area index; SER_max_, maximal rate of shoot expansion rate; TiR_max_, maximal value of tillering rate; LER_max_, maximum leaf elongation rate; The right side displays the Pearson correlation coefficients (r) and associated *P* value (blue: *T*. *durum*; red: *T. aestivum*).

### Plasticity of whole plant leaf area expansion under drought is driven by that of tiller appearance rate

Field conditions do not allow imposing well-defined water scenarios so we analyzed the sensitivity of leaf expansion processes to soil water potential in the platform only but with the insurance that the determined genetic variability could capture that observed in field conditions. Therefore, we analyzed the relative contributions of the sensitivity to SWP of the three underlying processes to that of whole-plant leaf expansion (calculated for all genotypes as in Fig. 3). In both species, the genetic variability of sensitivity of TiR to SWP was similar to that of SER (both were null at around −1 MPa; Fig. 3 and Fig.7). On the contrary, the sensitivity of LAR and LER to SWP was very low for both species (Fig. 5) and that of LAR was even negative for most genotypes (Fig.7).

**Figure 7.**
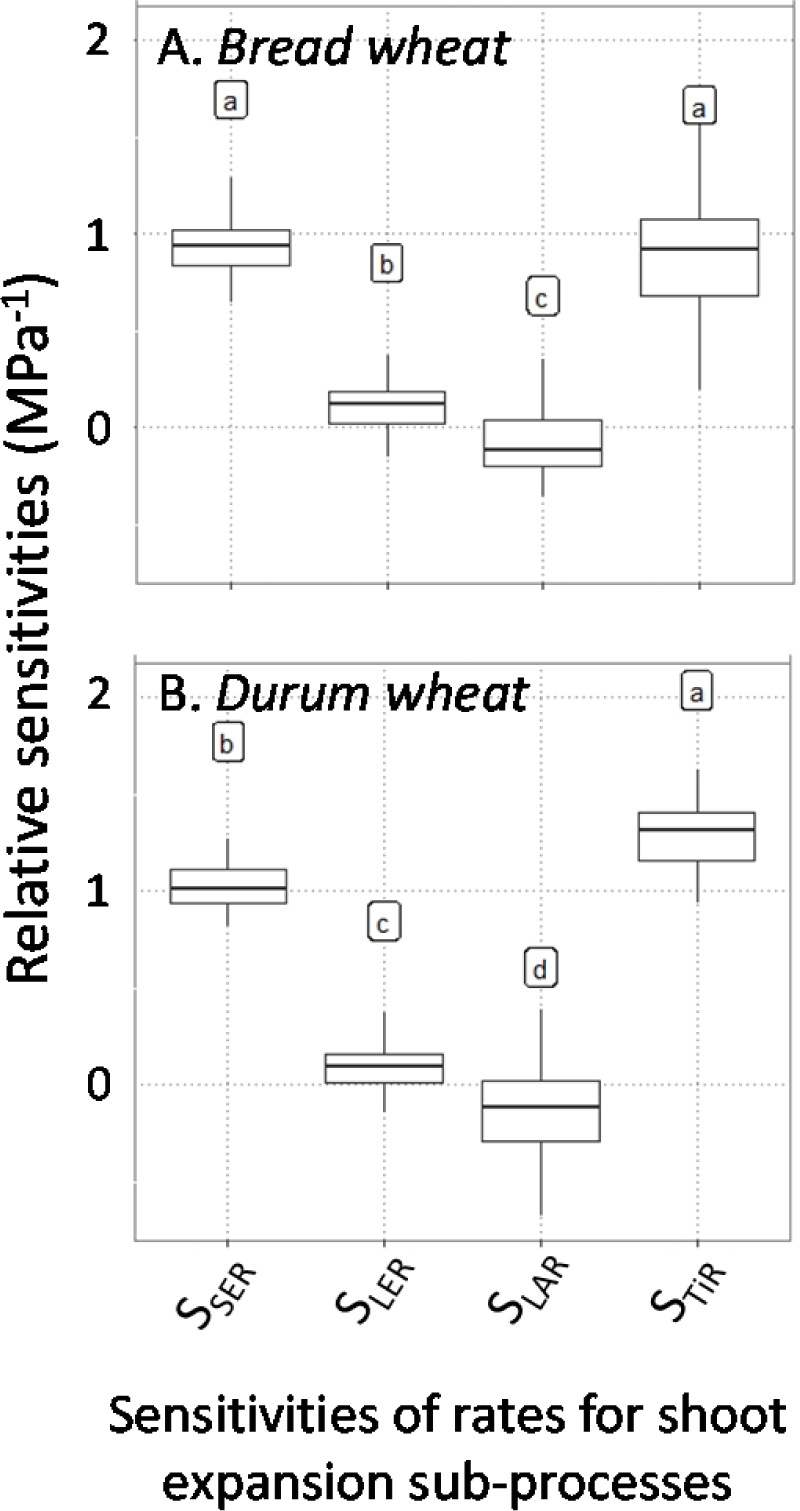
Box plot of sensitivities of shoot expansion rate to soil water potential (S_SER_), and that of its three underlying components, sensitivity of main-stem leaf appearance rate (*S*_LAR_), sensitivity of tiller appearance rate (*S*_TIR_), and sensitivity of leaf elongation rate (*S*_LER_) for 44 genotypes of bread wheat (A) and 38 genotypes of durum wheat (B). The letters indicate significant difference (Tukey test, *P* < 0.05) between variables. In each box plot, end of vertical lines represent from top to bottom, the 10^th^, and 90^th^ percentiles, horizontal lines represent from top to bottom, the 25^th^, 50^th^, and 75^th^ percentiles of the simulation.

An analysis of variance between *S*_SER_ and the three underlying sensitivities (Table 2B) suggested that *S*_SER_ was mainly driven by *S*_TiR_. Indeed, while the three underlying sensitivities were significantly associated with *S*_SER_, less variance was explained by *S*_LAR_ and *S*_LER_ than by *S*_TiR_. In both species, *S*_SER_ and *S*_TiR_ were significantly correlated (Fig. 8). However, it was not the case for *S*_LAR_ and *S*_LER_, which were not significantly correlated with *S*_SER_ for at least one species and when both species were considered together. As previously found in rice (Parent et al., 2010) or maize (Welcker et al., 2011), *S*_SER_ was negatively correlated with SER_max_. On the contrary, and unexpectedly, for both species *S*_LER_ and *S*_TiR_ were significantly correlated (Fig. 8), indicating strong links between the *S*_LER_ and *S*_TiR_.

**Figure 8.**
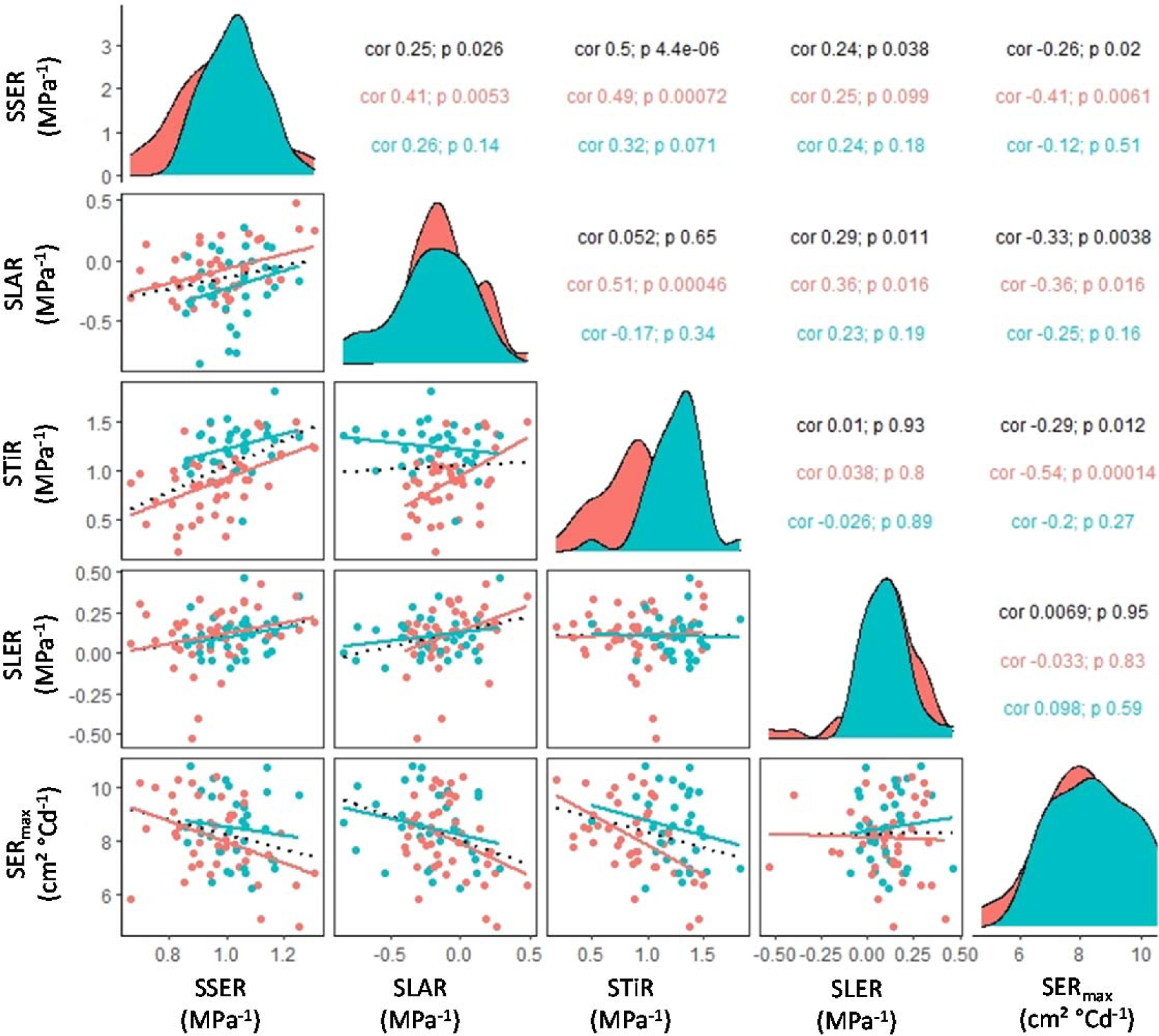
Scatter plot matrix of bivariate relationships between sensitivity of shoot expansion rate to soil water potential (S_SER_), its three underlying processes, sensitivity of leaf appearance rate (*S*_LAR_), sensitivity of tiller appearance rate (*S*_TiR_), sensitivity of leaf elongation rate (*S*_LER_) to soil water potential, and a potential covariable (maximum shoot expansion rate (SER_max_) for 38 genotypes of durum wheat (blue circles and lines) and 44 genotypes of bread wheat (red circles and lines) grown under well-watered conditions in the phenotyping platform. The distribution of each variable for each species is given in the diagonal. The upper half of the scatter plot matrix gives the Pearson’s correlation coefficients (*r*) and associated *P*-values, calculated for both species together (black font), and separately for durum wheat (blue font) and bread wheat (red font).

Overall, the above analyses showed that the sensitivity of SA to SWP was the emergent property of that of the three underlying processes, with *S*_TiR_ being the main contributor, followed by *S*_LER_ and *S*_LAR_.

## Discussion

This study aimed at determining the links between the underlying components of shoot expansion under drought and their relative contribution to the genetic variability of shoot expansion in wheat. By combining automated measurements in the platform and in the field together with large-scale manual measurements, we assembled a large dataset on leaf expansion processes at five scales, for 33 and 44 genotypes of durum wheat and bread wheat, respectively. We concluded that (i) growth traits measured in the platform at plant level can inform on canopy expansion in the field, with potentially large consequence on phenotyping protocols and variety evaluation. (ii) We simplified the scheme of the processes underlying the genetic variability of wheat growth rate, by quantifying the major impact of tillering under both optimal or drought conditions, the relative independence of the sensitivity of individual-leaf expansion compared to that of tillering and the very low sensitivity of leaf appearance rate. (iii) These results could have major impact on ecophysiological modeling of leaf growth rate under drought, by providing ranges for parameters values in the genetic diversity of bread wheat and durum wheat, specifying the links between parameters, simplifying the identification of ideal sets of genotypic parameters (ideotypes) adapted to contrasted drought scenarios.

### Growth traits measured in the platform at plant level can inform on canopy expansion in the field

Field phenotyping under drought faces several limitations (Langstroff et al., 2022): (i) the random and experimentally risky aspect of water deficit that may or may not occur under rainfed conditions; (ii) its very dynamic aspect, which complicates the analyses for plants with different phenology (Millet et al., 2019); (iii) The difficulty to quantify the water deficit according to soil depth and the associated root and soil/root conductance (Tardieu et al., 2017); (iv) The interaction of water deficit with other abiotic stresses occurring at the same time, in particular heat stress (Webber et al., 2022). It is therefore reasonable to focus the phenotyping of the responses to water-deficit in controlled-conditions. However, such platforms do not reflect agronomic conditions and a usual question is whether results obtained in controlled environment could be relevant to the field (Langstroff et al., 2022). Several results in this study indicate that growth traits measured in the platform on individual plants can inform on canopy expansion in the field:

-Under well-watered conditions, we showed that whole plant shoot expansion measured in the platform was significantly correlated with that of canopy expansion in the field. Less expected, the tillering rate measured in the platform was even more significantly associated with canopy expansion in the field.

-The ranking of genotypes based on the sensitivity of spike number in the field (Supplemental Fig S1) was also partly found on TiR and/or the growth of the whole plant. Indeed, the two genotypes of bread wheat of group A chosen to be the least sensitive in the field were also the least sensitive in the platform. The same was true for durum wheat.

This conclusion supported some previous studies showing that traits related to growth potential and growth sensitivity to water deficit can be conserved in the field and in the platform (Dignat et al., 2013; Parent et al., 2015). Although in the platform the number of tillers per plants was much higher than in the field, the relative responses of shoot growth processes to soil water deficit in the platform can be extrapolated to the field, or at least the ranking of genotypes found here can be considered as similar in the field and under controlled conditions. Such results can have in turn consequences in genetics, because a genetic analysis based on growth parameters should result in genetic markers of interest for field conditions, and modelling, because these parameters can be easily linked to genotypic parameters of crop models. It supports the idea that these highly-controlled platforms can have a place in academic research and pre-breeding pipelines, and that such place does not depend on the development (expected) of field phenotyping but rather on specific focus on adaptive traits such as drought response traits.

### Which sub-process structure the genetic variability of whole-plant leaf area expansion

Our results confirm our working hypothesis that the genetic variabilities of tiller appearance, leaf appearance and individual-leaf elongation rates could drive all together that of shoot expansion rate. In well-watered conditions, each of these three components was associated with the genetic diversity of shoot expansion. Their respective impact followed their genetic relative variability (here quantified by their coefficient of variation), with a higher proportion of the genetic diversity of shoot expansion explained by that of TiR, followed by LER and LAR. This was the case in both species, even in durum wheat which displayed a lower tillering rate in the platform compared to bread wheat (similar results in the field, Giunta et al., 2019). As expected, crop duration (here, only FLN was presented) participated to structure this genetic variability (Giunta et al., 2018). Final leaf number, as the main driver of crop cycle duration (in maize, Parent et al., 2018; and wheat, Jamieson et al., 1998) was associated with maximum value of growth processes (SER_max_, TiR_max_, LER_max_). In particular, TiR_max_ was positively associated with FLN, as predicted by the Fibonacci series that links leaf and tiller appearance (Bos and Neuteboom, 1998).

Under water deficit, we first showed that the sensitivity of leaf growth at the whole-plant level was a consequence of the sensitivities of its three components (all being significant in the ANOVA). However, most of the variance in *S*_SER_ was explained by *S*_TiR_. This was reflected by the relative sensitivities of all components compared to that of *S*_SER_ (Supplemental Fig. S7). Thus, water deficit did not alter the structure of the genetic variability of shoot expansion, with tiller appearance being the main explanatory process of this genetic diversity. As observed in other crops (Welcker et al., 2011; Parent et al., 2010), the potential growth was negatively correlated with the growth sensitivity to SWP. Again, this appeared to be mainly driven by the sensitivity of tillering, which was significantly negatively correlated with SER_max_.

### Consequences for crop improvement and ideotype studies

The set of constraints between traits defines the set of possibilities, informing breeders about the feasible combinations of traits (Leveau et al., 2021):

-In near-optimal conditions, an important issue for breeding is to develop a genotype with a high potential of growth and a low-sensitivity to drought (Welcker et al., 2022). This study shows that breaking this relationship between potential growth and sensitivity would not be straightforward in wheat for which we found strong correlations in both durum and bread wheat.

-In environments subject to severe terminal drought, a breeding strategy may be to favor varieties with limited shoot growth, to maintain a satisfactory soil water status at the end of the cycle during grain filling (Wasson et al., 2012). Our study shows that focusing on tillering to control shoot expansion can be a good strategy. First, we showed that TiR has proportionally larger genetic diversity than LAR or LER. Moreover, excess tillers are carbon sinks, of which at least the structural part will not contribute to grain yield (Berry et al., 2003). A selection strategy based on this high genetic variability and/or genes (Atsmon and Jacobs, 1977) or identified quantitative trait loci (Wang et al., 2016), seems therefore adequate for environments where the probability of having sufficient rainfall at the end of the cycle is low, thus not requiring any particular tillering plasticity, but instead a low constitutive number of tillers.

However, there are generally as many drought scenarios as sites / year combinations (Tardieu et al., 2018), so a preferable selection strategy would be to focus on adaptive processes and their plasticity (Reynolds and Langridge, 2016). Here, we have shown that tillering is the most plastic growth process determining whole-plant leaf expansion compared to other growth components, and is largely responsible for whole-plant growth plasticity under water deficit. Although somewhat less plastic, leaf elongation rate also appears to be a good target for modifying whole-plant leaf area plasticity.

Determining the set of such genotypic parameters of drought responses together with the sets of correlations between them can also simplify crop modelling and ideotyping studies (Senapati et al., 2022). Indeed, our study indicates that the responses of growth processes to soil water deficit cannot be considered as independent from cycle duration (here represented by FLN). This can have large consequence in *in-silico* studies. For example, for predicting the impact of climate change, one must consider the possible impact on such parameters if modifying the earliness of future genotypes. In ideotype studies, relationships between traits should be considered to prevent the finding of ideotypes which could not be achieved by breeding. Here, we described the structure of the genetic parameters of growth processes, suggesting a minimum set of genetic parameters needed for considering the genetic diversity of whole-plant leaf area expansion and it is plasticity.

## Material and methods

### Genetic material

In this study, we analyzed 44 bread wheat genotypes (*Triticum aestivum* L.) and 33 durum wheat genotypes (*Triticum turgidum* L. subsp. *durum* (Desf.) Husn.) that represented a large genetic variability within each species.

In bread wheat, 31 genotypes were selected from a genetic association panel of 195 lines sampled from a worldwide collection (Balfourier et al., 2019) by minimizing linkage disequilibrium and by imposing constraints on phenology and plant height (Paux et al., 2022). These 31 genotypes were selected to maximize the diversity of the responses of yield and spike number to water and nitrogen deficits (Supplemental Fig. S1A). In addition, four recombinant inbreed lines from the Multi-Reference Nested Association Mapping population (MR-NAM) were added for their contrasted transpiration efficiency (SUNTOP_73 and SUNTOP_90; Fletcher, 2020) or stay-green phenotype (SUNTOP_9 and SUNTOP_41; Christopher et al., 2021) together with their three parental lines (Suntop, Drysdale, Dharwar Dry). Three hybrids (SY_117079, HYB_14200387, and HTW_17000044) were also included together with their three parents (Akamar, Oregrain, and Sublim).

For durum wheat, the genotypes were selected from a panel of 253 genotypes composed of (i) 76 cultivars selected from the UNIBO durum wheat panel developed by the University of Bologna, Italy, (ii) 69 elite durum wheat cultivars selected from the panel described by Laido et al. (2013) and from the germplasm collection of the Council for Agricultural Research and Economics (CREA), (iii) 47 elite cultivars of the GPDUR panel developed by ARVALIS and INRAE, France, and (iv) 61 lines selected from 480 pre-breeding lines derived from an open-pollinated population based on the intercrossing of about 650 accessions from wild subspecies and elite lines (David et al., 2014). The durum wheat genotypes used in this study were selected to maximize the diversity of the responses of grain yield and grain number per square meter to water and nitrogen deficits (Supplemental Fig. S1B).

### Description of the phenotyping platform and experimental design

The experiment was carried out in the *PhenoArch* platform, hosted in the Montpellier Plant Phenotyping Platforms (MP3; https://www6.montpellier.inra.fr/lepse/M3P). The setup and the functioning of *PhenoArch* has been described in details eslewhere (Cabrera-Bosquet et al., 2016) but it has since been expanded to a capacity of 2,400 plants. Briefly, *PhenoArch* is a greenhouse equipped with conveyor belts carrying carts with one pot each (average plant density of 12.5 plant m^-2^). Each day, plants are transported towards either watering stations, allowing an accurate weighing of the pots and watering at target soil water potential (SWP), or imaging stations equipped with a top and a side RGB cameras. Micrometeorological conditions at the plant level are monitored every minute at ten positions in the glasshouse and average values are recorded every 15 min. Supplemental light was provided when outside solar radiation was below 300 W m^−2^ or to extend the photoperiod to 12 h. Thermal time was calculated from the appearance of leaf 3, which for all genotypes occurred after the transfer on the *PhenoArch* conveyor. Thermal time was calculated from hourly average meristem temperature using a beta function, with minimum, optimum and maximum temperatures of 0, 27.5, and 40 °C, respectively (Wang et al., 2017).

The dry weight of soil in each pot was calculated at the beginning of the experiment from the fresh weight of soil and soil water content determined in soil samples (three soil samples were taken every 60 pots). Soil water content was then determined by automatically weighing the pots and removing the weight of plants (estimated from plant imaging, see below) and the pot itself. A water-retention curve was previously obtained in several experiments (Prado et al., 2018) by fitting predawn leaf water potential of drying plants measured with a Scholander-type pressure chamber (Soil Moisture Equipment Corp., Santa Barbara, USA) against soil water content with a nonlinear equation (van Genuchten, 1980). This curve was validated with few measurements performed on additional plants and used to calculate the mean SWP in each pot at each weighing time.

Genotypes were separated into two groups. Group A (six replicates by genotype /water scenario combination) comprised four elite bread wheat cultivars and two elite durum wheat cultivars with contrasted behavior in the field in response to water deficit (Supplemental Fig. S1). Two bread wheat cultivars that appeared as “sensitive” (Zitnica and Couga), based their grain yield response under low-input conditions in the field and two “tolerant” (Albatros and Torril), with contrasted response of spike number per square meter. In the same way, we selected a “sensitive” durum wheat genotype (Nemesis), and a “tolerant” durum wheat (Lahan), based on the maintenance of grain number under drought with contrasted response of grain number per square meter (Supplemental Fig. S1B). From the six replicates of genotype / water scenario combinations, three were harvested at anthesis. Ten watering scenarios were tested for group A (see below) but only the four scenarios that were also tested in the group B were analyzed in this study. Group B (three replicates by genotype / water scenario combinations) comprised the other 43 bread wheat genotypes and 31 durum wheat genotypes (Nemesis was also in both groups).

### Plant growth conditions and drought scenarios in the phenotyping platform

Dry seeds were sown on 28 November 2019 at 2 cm depth in 3 x 5 cm Jiffy pellets (Jiffy Group, Oslo, Norway) and placed in a vernalization chamber where air temperature was controlled at 3.2 ± 0.5 °C with a PPFD of 25 µmol m^-2^ s^-1^ and 12 h photoperiod. After 7 weeks, seedlings were placed for 1 week under fluctuating conditions in the greenhouse (Supplemental Fig. S2). Seedlings were then transplanted together with their Jiffy pellet into 5-L plastic pots (0.17 m diameter, 0.24 m high) filled with a 30:70 (v:v) mixture of clay and organic compost. From that date, plants stayed on the *PhenoArch* greenhouse under fluctuating conditions (Supplemental Fig. S2) until final harvest at grain ripeness maturity on 12 May 2020.

### The four watering scenarios common to genotypes of group A and B analyzed in this study were

- *Scenario 1 (S1)*: well-watered conditions (control). SWP was maintained above −0.05 MPa during the whole plant growth cycle.
- *Scenario 2 (S2*): Late and short water deficit, a 3-weeks period of water deficit between the beginning of stem elongation (average of all genotypes) until anthesis of the earliest genotypes. Water was withheld on 12 February 2020 until SWP decreased to − 0.54 MPa, after which SWP was maintained at that value. At the end of the 3-weeks period of water deficit (6 March 2020), pots were rewatered over a 5-d period during which SWP was gradually increased to −0.25, −0.17, −0.125, −0.1, and −0.05 MPa, and then maintained above − 0.05 MPa until final harvest
- *Scenario 3 (S3)*: Early and short water deficit, a 3-weeks period of water deficit starting at the beginning of tillering when the plants had three to four visible leaves depending on the genotype. Water was withheld on 3 February 2020 until SWP decreased to − 0.54 MPa. At the end of the 3-weeks period (28 February 2020), pots were slowly rewatered as in S2.
- *Scenario 4 (S4)*: Early and long water deficit, a 31-d period of water deficit starting at the beginning of tillering (average of all genotypes, 3 February 2020) until anthesis of the earliest genotypes (6 March 2020).

### Automatic measurements of whole plant shoot area in the phenotyping platform

An analysis pipeline was used to calculate whole plant leaf area (shoot SA) and fresh weight from RGB images (2056 × 2454 px) taken from 13 angle views (12 side views and one top view) every 2 to 3 d for each plant in the platform. Briefly, green pixels from each image were segmented from those of the background and projected plant area were then calculated using calibration curves obtained previously using reference objects to convert pixels into projected plant green area (Brichet et al., 2017). Multiple-linear regression models (parametrized from ground-truth measurements from this experiment together with previous experiments) were then used to calculate whole plant shoot area and plant fresh weight.

### Determination of the number of appeared main-stem leaf and green tiller number per plant in the phenotyping platform

The number of leaves on the main-stem (LN) and green tillers per plant (TN) of each plant were manually counted twice a week from the beginning of tillering to anthesis. The former was calculated as Haun (1973):

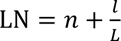

where *n* is the number of mature leaves (with the ligule appeared), *l* is the exposed length of leaf *n*+1 at the time of measurement, and *L* is the final length of the blade of leaf *n*+1. The exposed length of a leaf was measured with a ruler as the distance from leaf tip to the upper collar of the sheath tube.

Final (mature) leaf length and width of each newly mature leaf on the main stem were measured with a ruler. Cumulative leaf length (CLL) was calculated by summing the length of all appeared leaves.

### Calculation of rates and sensitivities to soil water deficit in the phenotyping platform

We analyzed four processes (Table 1), the rate of whole plant leaf area expansion (SER) and the three underlying processes, the rates of tiller appearance (TiR), leaf appearance (LAR), and individual leaf elongation on the main stem (LER). For each process, several variables were compared between genotypes (Table 1): (i) the time course of the corresponding variable as a function of thermal time; (ii) its maximum value; (iii) the time course of the related rate as a function of thermal time; (iv) its maximum (for SER, TiR, and LER) or average value (for LAR); and (v) the relative sensitivity of the rate to SWP (i.e. the slope of the relationship between relative rate and SWP).

SA and TN versus thermal time were fitted to a cubic smoothing spline function equation using the smooth.spline() function of the R package *stats* (R Core Team, 2018) with *spar* parameter set at 0.6 and 0.4, respectively. LN versus thermal time was fitted to a linear model forced to 3 at 0°Cd (thermal time started at leaf 3 emergence). For the control treatment (scenario 1), the spline function or linear model was fitted to the data of all the plants of each genotype (*n* = 3 and 6 replicates for group B and A, respectively), in order to define a reference on which calculating relative rate in water deficit. For the water deficit scenario, the data for each plant were fitted separately. The rate of each process was then calculated as the first derivative of the fitted spline or linear regression. We calculated LAR as the slope of the linear regression and LER was calculated as CLL x LAR_mean_. Finally, for each process and each plant subjected to water deficit, relative rates were calculated by dividing the absolute rates by those of well-watered plants (average of all plants of the same genotype).

For each process and genotype, we defined a period during which the rate of the process for the well-watered planted was high enough to accurately calculate SWP sensitivity. This period was defined from the time the rate was at 30% of its maximum value and either the time it had decreased to 20% of its maximum value or the date of the first rewatering, whichever occurred first (Fig. 1B,D). For each genotype, the sensitivity of each process to soil water deficit was calculated during this period as the slope of the relationship between relative rate versus SWP forced to 1 when SWP is null (Fig. 1D).

### Description of field trials

Two field trials were carried out at the ARVALIS experimental station at Gréoux-les-Bains, France (43°44’ N, 5°51’ E, 293 m a.s.l.) during the 2017-2018 and 2018-2019 growing seasons with a subset of 33 durum wheat (Supplemental Table. S1) and 44 bread wheat cultivars (Supplemental Table. S2), respectively. Crops were sown on 29 October 2017 and 20 November 2018 at a density of 300 seeds m^-2^. Plots consisted in 11 rows with an inter-row spacing of 0.175 m and a surface area of 11.9 m^-2^. In each trial, cultivars were randomized in each of the two blocks of replicates. Soil water content at 30− cm and 60-cm below the soil surface was monitored throughout the growing season using tensiometric probes. All crop inputs including weed, disease and pest control, irrigation, and nitrogen, potassium, phosphate, and sulfur fertilizers, replicated local practices to prevent N, non-N nutrients, weeds, diseases and pests from limiting grain yield. The bread and durum wheat trials received a total of 70 and 200 mm of irrigation, applied in two and six applications between growth stages (Zadoks et al., 1974) first node detected and medium milk, and 180 and 230 kg N ha^-1^ of fertilizer, applied as ammonium nitrate in three and four splits between growth stages three leaves emerged and heading complete, respectively.

### Calculation of canopy expansion rates in the field

Green area index (GAI) was determined with a self-guided rover equipped with RGB cameras mounted at 0° and 45° zenithal view angles (PHENOMOBILE V1; Madec et al., 2017) using the procedures described by (Madec et al., 2017). In both trials, measurements were taken every about 500°Cd during the whole season, and a higher frequency (every 100°Cd) from 1000°Cd and 1500°Cd after sowing. The rates of change of GAI were calculated as the first derivative of a cubic smoothing spline fitted to the absolute values versus thermal time after sowing. The best *spar* parameter of the R function *smooth.spline*() was first estimated for each genotype / treatment combination and the median value (0.5) was then used for all the plots.

### Other statistical analyses

All data analyses were performed with the R statistical software program R (R Core Team, 2018). The association between SER_max_ with LAR_mean_, LER_max_, TIR_max_, and FLN as predictor was analyzed with an ANOVA. In the same way, an anova was used to test the association between the sensitivity of shoot expansion rate with *S*_LAR_, *S*_TIR_, and *S*_LER_.

## Author contributions

BP, PM and FG designed the research and the experiments; FG, SL, BP, LCB, NL and PM performed the experiment in the phenotyping platform; KB defined and supervised the field experiments; SL, BP, and PM analyzed the data; BP wrote the first draft of the manuscript; all authors contributed to the revision of the manuscript.

## Acknowledgements

We gratefully acknowledge the skilled technical help of Francesco Cadeddu, Romane Le Roy, Francesca Mureddu, Alissar Nehmed, and Benoit Suard in the *PhenoArch* plateform, and Magali Camous and Olivier Moulin in the field. We also thank Prof. Dr. Xavier Draye (UCL, Belgium), Dr. Jacques Le Gouis (UMR GDEC, INRAE, Clermont-Ferrand, France), Dr. Pierre Roumet (AGAP Institut, INRAE, Montpellier, France), Dr. Nicola Pecchioni (CREA, Foggia, Italy) for their contribution to the selection of the genotypes used in this study, and Dr. Pierre Roumet, Dr. Jacques Le Gouis, and Dr. Karine Chenu (University of Queensland, Australia), Dr. Yan Manes (Syngenta, France) for providing the seeds of the genotypes used in this study.

## Supplemental Material

The following materials are available in the online version of this article.

**Supplemental Table S1.** List of the bread wheat genotypes used in this study.

**Supplemental Table S2.** List of the durum wheat genotypes used in this study.

**Supplemental Table S3.** Genotypic means of all variables analyzed in the platform for bread wheat genotypes.

**Supplemental Table S4.** Genotypic means of all variables analyzed in the platform for durum wheat genotypes.

**Supplemental Figure S1**. Relationship between the weighted sensitivity rank for grain yield and spike number per square meter for the panel of 210 bread wheat genotypes.

**Supplemental Figure S2**. Hourly air temperature, relative humidity and vapor pressure deficit, and photosynthetically photon flux density during the experiment in the high-throughput plant phenotyping platform *PhenoArch*.

**Supplemental Figure S3**. Time courses of tiller appearance and sensitivity of tiller appearance rate to soil water potential for the four bread wheat and two durum wheat cultivars of group A under the four watering scenarios.

**Supplemental Figure S4**. Time courses of cumulative leaf length per plant and sensitivity of leaf elongate rate to soil water potential for the four bread wheat and two durum wheat cultivars of group A under the four watering scenarios.

**Supplemental Figure S5**. Time courses of main-stem leaf number and sensitivity of leaf appearance rate to soil water potential for the four bread wheat and two durum wheat cultivars of group A under the four watering scenarios.

**Supplemental Figure S6**. Scatter plot matrix of bivariate relationships between maximum shoot expansion rate, its three underlying processes and two potential covariables.

**Supplemental Figure S7**. Relative sensitivities of shoot expansion rate, main-stem leaf appearance rate, main-stem leaf elongation rate, and tiller appearance rate to soil water potential for the 44 bread wheat and the 33 durum wheat genotypes studied.

## References

1. Atsmon D, Jacobs E (1977) A Newly Bred ‘Gigas’ Form of Bread Wheat (Triticum aestivum L.): Morphological Features and Thermo-photoperiodic Responses1. Crop Science 17: cropsci1977.0011183X001700010010x

2. Balfourier F, Bouchet S, Robert S, De Oliveira R, Rimbert H, Kitt J, Choulet F, Appels R, Feuillet C, Keller B, Praud S, Baumann U, Budak H, Rogers J, Eversole K, Alaux M, Bejar B, Lafarge S, Lagendijk E, Derory J, Le Gouis J, Paux E (2019) Worldwide phylogeography and history of wheat genetic diversity. Science Advances 5: eaav0536

3. Baumont M, Parent B, Manceau L, Brown HE, Driever SM, Muller B, Martre P (2019) Experimental and modeling evidence of carbon limitation of leaf appearance rate for spring and winter wheat. Journal of Experimental Botany 70: 2449–2462

4. Beauchêne K, Leroy F, Fournier A, Huet C, Bonnefoy M, Lorgeou J, de Solan B, Piquemal B, Thomas S, Cohan J-P (2019) Management and Characterization of Abiotic Stress via PhénoField®, a High-Throughput Field Phenotyping Platform. Frontiers in Plant Science 10: 904

5. Belaygue C, Wery J, Cowan A, Tardieu F (1996) Contribution of Leaf Expansion, Rate of Leaf Appearance, and Stolon Branching to Growth of Plant Leaf Area under Water Deficit in White Clover. Crop Science 36: cropsci1996.0011183X003600050028x

6. Berry PM, Spink JH, Foulkes MJ, Wade A (2003) Quantifying the contributions and losses of dry matter from non-surviving shoots in four cultivars of winter wheat. Field Crops Research 80: 111–121

7. Bos HJ, Neuteboom JH (1998) Growth of individual leaves of spring wheat (Triticum aestivum L.) as influenced by temperature and light intensity. Annals of Botany 81: 141–149

8. Brichet N, Fournier C, Turc O, Strauss O, Artzet S, Pradal C, Welcker C, Tardieu F, Cabrera-Bosquet L (2017) A robot-assisted imaging pipeline for tracking the growths of maize ear and silks in a high-throughput phenotyping platform. Plant Methods 13: 96

9. Cabrera-Bosquet L, Fournier C, Brichet N, Welcker C, Suard B, Tardieu F (2016) High-throughput estimation of incident light, light interception and radiation-use efficiency of thousands of plants in a phenotyping platform. New Phytologist 212: 269–281

10. Christopher M, Paccapelo V, Kelly A, Macdonald B, Hickey L, Richard C, Verbyla A, Chenu K, Borrell A, Amin A, Christopher J (2021) QTL identified for stay-green in a multi-reference nested association mapping population of wheat exhibit context dependent expression and parent-specific alleles. Field Crops Research 270: 108181

11. Correia PMP, Westergaard JC, da Silva AB, Roitsch T, Carmo-Silva E, da Silva JM (2022) High-throughput phenotyping of physiological traits for wheat resilience to high temperature and drought stress. Journal of Experimental Botany 73: 5235–5251

12. David J, Holtz Y, Ranwez V, Santoni S, Sarah G, Ardisson M, Poux G, Choulet F, Genthon C, Roumet P, Tavaud-Pirra M (2014) Genotyping by sequencing transcriptomes in an evolutionary pre-breeding durum wheat population. Molecular Breeding 34: 1531–1548

13. Dhakar R, Nagar S, Sehgal VK, Jha PK, Singh MP, Chakraborty D, Mukherjee J, Prasad PVV (2023) Balancing water and radiation productivity suggests a clue for improving yields in wheat under combined water deficit and terminal heat stress. Frontiers in Plant Science 14: 14

14. Dignat G, Welcker C, Sawkins M, Ribaut JM, Tardieu F (2013) The growths of leaves, shoots, roots and reproductive organs partly share their genetic control in maize plants. Plant, Cell & Environment 36: 1105–1119

15. Faralli M, Williams KS, Han J, Corke FMK, Doonan JH, Kettlewell PS (2019) Water-Saving Traits Can Protect Wheat Grain Number Under Progressive Soil Drying at the Meiotic Stage: A Phenotyping Approach. Journal of Plant Growth Regulation 38: 1562-1573

15. Fletcher AL (2020) Understanding the genetic and physiological basis of transpiration efficiency in Australian wheat. PhD Thesis. The University of Queensland

16. Giunta F, De Vita P, Mastrangelo AM, Sanna G, Motzo R (2018) Environmental and Genetic Variation for Yield-Related Traits of Durum Wheat as Affected by Development. Frontiers in Plant Science 9

17. Giunta F, Pruneddu G, Zuddas M, Motzo R (2019) Bread and durum wheat: Intra- and inter-specific variation in grain yield and protein concentration of modern Italian cultivars. European Journal of Agronomy 105: 119–128

18. Hatfield JL, Dold C (2019) Water-Use Efficiency: Advances and Challenges in a Changing Climate. Frontiers in Plant Science 10

19. Haun JR (1973) Visual Quantification of Wheat Development1. Agronomy Journal 65: 116

20. Hendriks W, Kirkegaard JA, Lilley JM, Gregory PJ, Rebetzke GJ (2016) A tillering inhibition gene influences root-shoot carbon partitioning and pattern of water use to improve wheat productivity in rainfed environments. Journal of Experimental Botany 67: 327–340

21. Jamieson PD, Brooking IR, Semenov MA, Porter JR (1998) Making sense of wheat development: a critique of methodology. Field Crops Research 55: 117-127

22. Janni M, Pieruschka R (2022) Plant phenotyping for a sustainable future. Journal of Experimental Botany 73: 5085–5088

23. Koch G, Rolland G, Dauzat M, Bédiée A, Baldazzi V, Bertin N, Guédon Y, Granier C (2019) Leaf Production and Expansion: A Generalized Response to Drought Stresses from Cells to Whole Leaf Biomass—A Case Study in the Tomato Compound Leaf. Plants 8: 409

24. Lacube S, Manceau L, Welcker C, Millet EJ, Gouesnard B, Palaffre C, Ribaut J-M, Hammer G, Parent B, Tardieu F (2020) Simulating the effect of flowering time on maize individual leaf area in contrasting environmental scenarios. Journal of Experimental Botany 71: 5577–5588

25. Laido G, Mangini G, Taranto F, Gadaleta A, Blanco A, Cattivelli L, Marone D, Mastrangelo AM, Papa R, De Vita P (2013) Genetic Diversity and Population Structure of Tetraploid Wheats (Triticum turgidum L.) Estimated by SSR, DArT and Pedigree Data. Plos One 8

26. Langstroff A, Heuermann MC, Stahl A, Junker A (2022) Opportunities and limits of controlled-environment plant phenotyping for climate response traits. Theoretical and Applied Genetics 135: 1-16

27. Leveau S, Parent B, Zaka S, Martre P (2021) Differential sensitivity to temperature and evaporative demand in wheat relatives. Journal of Experimental Botany

28. Madec S, Baret F, de Solan B, Thomas S, Dutartre D, Jezequel S, Hemmerlé M, Colombeau G, Comar A (2017) High-Throughput Phenotyping of Plant Height: Comparing Unmanned Aerial Vehicles and Ground LiDAR Estimates. Frontiers in Plant Science 8

29. Millet EJ, Kruijer W, Coupel-Ledru A, Prado SA, Cabrera-Bosquet L, Lacube S, Charcosset A, Welcker C, van Eeuwijk F, Tardieu F (2019) Genomic prediction of maize yield across European environmental conditions. Nature Genetics 51: 952-+

30. Parent B, Leclere M, Lacube S, Semenov MA, Welcker C, Martre P, Tardieu F (2018) Maize yields over Europe may increase in spite of climate change, with an appropriate use of the genetic variability of flowering time. Proceedings of the National Academy of Sciences: 201720716

31. Parent B, Shahinnia F, Maphosa L, Berger B, Rabie H, Chalmers K, Kovalchuk A, Langridge P, Fleury D (2015) Combining field performance with controlled environment plant imaging to identify the genetic control of growth and transpiration underlying yield response to water-deficit stress in wheat. Journal of Experimental Botany 66: 5481-5492

32. Paux E, Lafarge S, Balfourier F, Derory J, Charmet G, Alaux M, Perchet G, Bondoux M, Baret F, Barillot R, Ravel C, Sourdille P, Le Gouis J, Consortium obotB (2022) Breeding for Economically and Environmentally Sustainable Wheat Varieties: An Integrated Approach from Genomics to Selection. Biology 11: 149

33. Pierik R, Fankhauser C, Strader LC, Sinha N (2021) Architecture and plasticity: optimizing plant performance in dynamic environments. Plant Physiology 187: 1029-1032

34. Prado SA, Cabrera-Bosquet L, Grau A, Coupel-Ledru A, Millet EJ, Welcker C, Tardieu F (2018) Phenomics allows identification of genomic regions affecting maize stomatal conductance with conditional effects of water deficit and evaporative demand. Plant Cell and Environment 41: 314-326

35. R Core Team (2018) R: A Language and Environment for Statistical Computing. R Foundation for Statistical Computing, Vienna, Austria

36. Reynolds M, Langridge P (2016) Physiological breeding. Current Opinion in Plant Biology 31: 162-171

37. Richards RA (1988) A TILLER INHIBITOR GENE IN WHEAT AND ITS EFFECT ON PLANT-GROWTH. Australian Journal of Agricultural Research 39: 749–757

38. Rymen B, Sugimoto K (2012) Tuning growth to the environmental demands. Current Opinion in Plant Biology 15: 683–690

39. Sanad MNME, Campbell KG, Gill KS (2016) Developmental program impacts phenological plasticity of spring wheat under drought. Botanical Studies 57: 35

40. Senapati N, Semenov MA, Halford NG, Hawkesford MJ, Asseng S, Cooper M, Ewert F, van Ittersum MK, Martre P, Olesen JE, Reynolds M, Rotter RP, Webber H (2022) Global wheat production could benefit from closing the genetic yield gap. Nature Food 3: 532–541

41. Suneja Y, Gupta AK, Bains NS (2019) Stress Adaptive Plasticity: Aegilops tauschii and Triticum dicoccoides as Potential Donors of Drought Associated Morpho-Physiological Traits in Wheat. Frontiers in Plant Science 10

42. Tardieu F, Cabrera-Bosquet L, Pridmore T, Bennett M (2017) Plant Phenomics, From Sensors to Knowledge. Curr Biol 27: R770–R783

43. Tardieu F, Draye X, Javaux M (2017) Root Water Uptake and Ideotypes of the Root System: Whole-Plant Controls Matter. Vadose Zone Journal 16

44. Tardieu F, Simonneau T, Muller B (2018) The Physiological Basis of Drought Tolerance in Crop Plants: A Scenario-Dependent Probabilistic Approach. *In* SS Merchant, ed, Annual Review of Plant Biology, Vol 69, Vol 69, pp 733–759

45. van Genuchten MT (1980) A Closed-form Equation for Predicting the Hydraulic Conductivity of Unsaturated Soils. Soil Science Society of America Journal 44: 892–898

46. Verbraeken L, Wuyts N, Mertens S, Cannoot B, Maleux K, Demuynck K, De Block J, Merchie J, Dhondt S, Bonaventure G, Crafts-Brandner S, Vogel J, Bruce W, Inzé D, Maere S, Nelissen H (2021) Drought affects the rate and duration of organ growth but not inter-organ growth coordination. Plant Physiology 186: 1336–1353

47. Wang E, Martre P, Zhao ZG, Ewert F, Maiorano A, Rotter RP, Kimball BA, Ottman MJ, Wall GW, White JW, Reynolds MP, Alderman PD, Aggarwal PK, Anothai J, Basso B, Biernath C, Cammarano D, Challinor AJ, De Sanctis G, Doltra J, Fereres E, Garcia-Vila M, Gayler S, Hoogenboom G, Hunt LA, Izaurralde RC, Jabloun M, Jones CD, Kersebaum KC, Koehler AK, Liu LL, Muller C, Kumar SN, Nendel C, O’Leary G, Olesen JE, Palosuo T, Priesack E, Rezaei EE, Ripoche D, Ruane AC, Semenov MA, Shcherbak I, Stockle C, Stratonovitch P, Streck T, Supit I, Tao FL, Thorburn P, Waha K, Wallach D, Wang ZM, Wolf J, Zhu Y, Asseng S (2017) The uncertainty of crop yield projections is reduced by improved temperature response functions. Nature Plants 3

48. Wang ZQ, Liu YX, Shi HR, Mo HJ, Wu FK, Lin Y, Gao S, Wang JR, Wei YM, Liu CJ, Zheng YL (2016) Identification and validation of novel low-tiller number QTL in common wheat. Theoretical and Applied Genetics 129: 603–612

49. Wasson AP, Richards RA, Chatrath R, Misra SC, Prasad SVS, Rebetzke GJ, Kirkegaard JA, Christopher J, Watt M (2012) Traits and selection strategies to improve root systems and water uptake in water-limited wheat crops. Journal of Experimental Botany 63: 3485–3498

50. Webber H, Rezaei EE, Ryo M, Ewert F (2022) Framework to guide modeling single and multiple abiotic stresses in arable crops. Agriculture Ecosystems & Environment 340

51. Welcker C, Sadok W, Dignat G, Renault M, Salvi S, Charcosset A, Tardieu F (2011) A Common Genetic Determinism for Sensitivities to Soil Water Deficit and Evaporative Demand: Meta-Analysis of Quantitative Trait Loci and Introgression Lines of Maize. Plant Physiology 157: 718–729

52. Welcker C, Spencer NA, Turc O, Granato I, Chapuis R, Madur D, Beauchene K, Gouesnard B, Draye X, Palaffre C, Lorgeou J, Melkior S, Guillaume C, Presterl T, Murigneux A, Wisser RJ, Millet EJ, van Eeuwijk F, Charcosset A, Tardieu F (2022) Physiological adaptive traits are a potential allele reservoir for maize genetic progress under challenging conditions. Nature Communications 13

53. Zadoks JC, Chang TT, Konzak CF (1974) Decimal code for growth stages of cereals. Weed Research 14: 415–421

